# Grb7 knockout mice develop normally but litters born to knockout females fail to thrive

**DOI:** 10.1101/2023.06.29.546912

**Authors:** Kristopher A. Lofgren, Paraic A. Kenny

## Abstract

Growth factor receptor-bound 7 (Grb7) is an adaptor protein involved in signal transduction downstream of multiple receptor tyrosine kinases, including ERBB, FGFR and PDGFR pathways. Experimental studies have implicated Grb7 in regulating cell proliferation, survival, migration and invasion through its large repertoire of protein-protein interactions. Here, we describe the generation and characterization of a *Grb7* knockout mouse. These mice are viable and fertile. A *lacZ* knock-in reporter was used to visualize *Grb7* promoter activity patterns in adult tissues, indicating widespread Grb7 expression in glandular epithelium, the central nervous system and other tissues. The sole defect observed in these animals was a failure of *Grb7* knockout females to successfully raise pups to weaning age, a phenotype that was independent of both paternal and pup genotypes. These data suggest a regulatory role for Grb7 in mammary lactational physiology.

## Introduction

Growth factor receptor-bound 7 (Grb7) is a member of the Grb7/Grb10/Grb14 family of adapter proteins. The human (532 amino acid) and mouse (535 amino acid) Grb7 proteins have the same multidomain structure and are 93% homologous. Grb7 consists of several domains facilitating protein-protein interactions and lacks enzymatic activity. From the N to C terminus, domains include a proline-rich domain (PR), a Ras-associating (RA) domain, a pleckstrin homology (PH) domain, a phosphotyrosine interacting region (PIR), and a Src homology 2 (SH2) domain.

Previously published studies have demonstrated *Grb7* expression in a variety of normal tissues, however the list is not comprehensive. Margolis et al. (Margolis et al., 1992) cloned *Grb7* and detected its expression in murine liver, kidney, and gonad tissue by northern blot. Leavey et al. (Leavey et al., 1998) detailed expression in the embryonic gut, lung, and in the embryonic and adult kidney. Frantz et al (Frantz et al., 1997) identified *Grb7* transcript in the pancreas, with lesser amounts in the kidney, prostate, small intestine, and placenta. While these prior studies using RNA analysis of bulk tissues have provided general information on the distribution of *Grb7* in organs, granular detail on the tissue compartments expressing *Grb7* is not comprehensive.

Additionally, several studies demonstrated *GRB7* alterations in various human cancers suggesting a role in malignancy. *GRB7* was first shown to be co-amplified with *ERBB2* in breast cancer (Stein et al., 1994), but alterations are also reported in esophageal (Tanaka et al., 1997), gastric (Kishi et al., 1997), testicular germ cell (McIntyre et al., 2005), pancreatic (Tanaka et al., 2006), hepatocellular (Itoh et al., 2007), ovarian (Wang et al., 2010), colorectal (Huang et al., 2012), thyroid (Tang et al., 2020), and bladder (Zheng et al., 2020) cancers. Peptide (Tanaka et al., 2006; Watson et al., 2017) and small molecule (Ambaye et al., 2011) inhibitors of GRB7 have been developed and have anticancer activity in pre-clinical models (Giricz et al., 2012; Pero et al., 2007) suggesting that GRB7 may be functionally important in these malignancies.

Using various experimental approaches, Grb7 has been proposed to contribute to regulation of cell proliferation, survival, migration, and invasion through its interacting proteins, which include EGFR/ErbB2/ErbB3, Shc, Ret, PDGFR, FGFRs, EphB1, c-Kit, FAK, Tek/Tie2, insulin receptor, caveolin, and calmodulin (reviewed in Chu et al., 2019; Han et al., 2001). Even though Grb7 has been implicated in many cellular processes, the extent to which Grb7 is required for normal development and physiology is unclear. Here we describe the characterization of mice with *Grb7* deletion and also take advantage of a *lacZ* knock-in allele to generate an atlas of *in vivo Grb7* expression in the adult mouse.

## Methods

### Mouse strains and breeding

All animal use was approved by the UW-La Crosse Institutional Animal Care and Use Committee and practices in compliance with the NIH Guide for the Care and Use of Laboratory Animals. Mice were housed in individually ventilated, specific pathogen-free caging on a standard 12 hr light/ 12 hr dark cycle. Grb7 heterozygous crosses were used to expand the colony.

*Grb7* “knockout-first” founders were purchased from MMRRC (*Grb7*^*tm1a(EUCOMM)Wtsi*^, MMRRC:041170-UCD, MGI:4453768, C57BL/6N background), expanded in-house, and genotyped with the following primers: G7fwd (T+) GGATTGGCATTTTGTCTG; G7rev (B-) AAAGCCAGTGTTCAGCCTCC; CAS_R1 rev (T-) TCGTGGTATCGTTATGCGCC. Mice expressing β-actin driven flp recombinase (Rodriguez et al., 2000) were purchased from The Jackson Laboratory (Stock 005703, B6.Cg-Tg(ACTFLPe)9205Dym/J, MGI:2448985). Beta-actin FLPe genotyping primers used to determine FLP transmission are Fwd: GTCCACTCCCAGGTCCAACTGCAGCCCAAG; Rev: CGCTAAAGAAGTATATGTGCCTACTAACGC. Cre-deleter mice expressing ubiquitous CMV-driven Cre recombinase (Schwenk et al., 1995) were purchased from The Jackson Laboratory (Stock 006054, B6.C-Tg(CMV-cre)1Cgn/J, MGI:2176180). Primers used for genotyping this strain are as follows: Fwd: GCGGTCTGGCAGTAAAAACTATC; Rev: GTGAAACAGCATTGCTGTCACTT

Primers used to confirm *Grb7* allele recombination events (and their IDs in figure 1B) are as follows:

**Figure 1.**
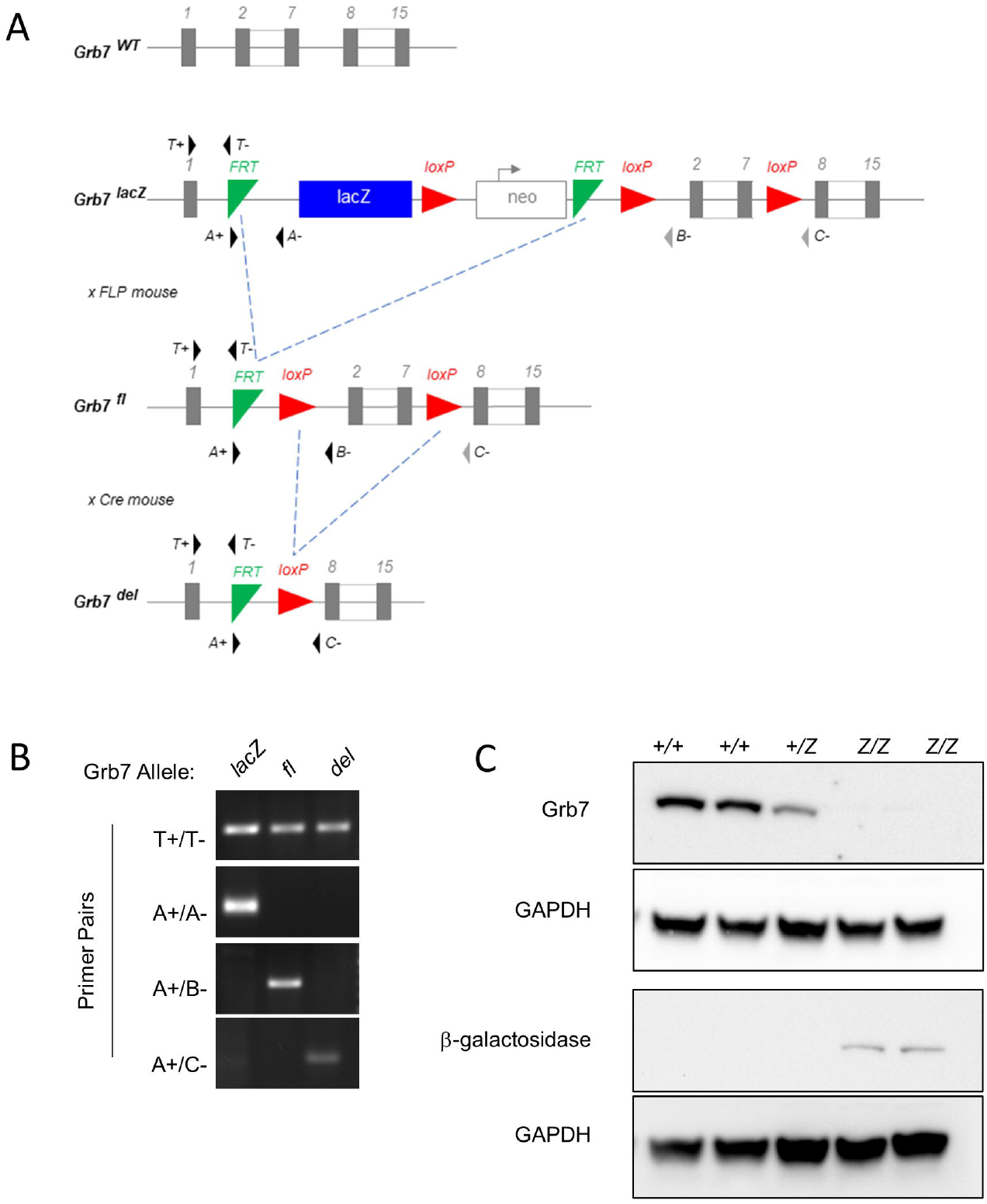
Generation of a *Grb7* knockout mouse. **A**. Allele descriptions, primer locations (arrowheads), and breeding schema. The wild-type *Grb7* allele is shown for reference (top), with variations below. Mice with the “knockout-first” *lacZ* gene-trap allele (*Grb7*^*lacZ*^) were bred to mice carrying an actin-driven *FLP* recombinase allele, resulting in excision of the *lacZ* cassette, yielding a conditional knockout allele, *Grb7*^*FL*^. A subsequent cross with mice expressing CMV-driven *cre* recombinase removed *Grb7* exons 2-7, yielding the knockout allele (*Grb7*^*del*^). **B**. PCR verification from tail biopsy DNA of stepwise generation of each allele: common primer set for transgenic allele (T+/T-), *Grb7*^*lacZ*^ allele (A+/A-), *Grb7*^*FL*^ (A+/B-) and *Grb7*^*-*^ (A+/C-primers). **C**. Western blotting of total liver lysate from *Grb7*^*+/+*^, *Grb7*^*+/lacZ*^, and *Grb7*^*lacZ/lacZ*^ mice for Grb7 and β-galactosidase.

**T+** G7 fwd: TGGGATTGGCATTTTGTCTG

**T-** CAS_R1 rev: TCGTGGTATCGTTATGCGCC

Tm1c Fwd (**A+**): AAGGCGCATAACGATACCAC

5’FRT Rev **(A-)**: CCACAACGGGTTCTTCTGTT

G7 Rev (**B-**): AAAGCCAGTGTTCAGCCTCC

Floxed LRev (**C-**): ACTGATGGCGAGCTCAGACC

### Knockout Mouse Phenotyping

Healthy adult *Grb7* wild-type, heterozygous and knockout littermates were submitted to the Comparative Pathology Laboratory (Research Animal Resources Center, University of Wisconsin, Madison, WI). Submitted females were aged 9 weeks and males were 11 weeks. Mice were monitored for abnormal activity prior to euthanasia and externally examined after euthanasia. A gross necropsy was performed with histopathological follow-up. Analysis was performed by veterinary pathologists on staff at the UW School of Veterinary Medicine.

### Tissue Harvesting and Processing

Mice were humanely euthanized with carbon dioxide and dissected. Tissues being harvested for formalin fixation and paraffin embedding were rinsed with PBS and placed in 10% neutral buffered formalin for a minimum of 48 hours at room temperature. Bony specimens were decalcified (Shandon TBD-1 decalcifier, Thermo Scientific) prior to further processing. Samples were grossed and trimmed if necessary, placed in histology cassettes and processed through graded alcohols, xylene, and paraffin in an overnight protocol on a Leica STP120 automated tissue processor.

For protein isolation, tissues were dissected and snap frozen in liquid nitrogen. Tissue pieces were thawed in CellLytic M lysis buffer (Sigma C2978) and homogenized with a motorized mortar and pestle. Samples were centrifuged at 10,000x g for 10 minutes in a refrigerated centrifuge.

### Western Blotting

Whole cell lysates were quantified by BCA, loaded on a 4-20% polyacrylamide gel, and then subjected to SDS-PAGE and transferred to PVDF. Membranes were blocked in 5% milk-PBST and incubated with 1% milk-PBST containing rabbit anti-Grb7 (C-20, Santa Cruz sc606), anti-β-galactosidase (Invitrogen A11132), or anti-GAPDH (Cell Signaling 2118) overnight at 4°C with constant rocking. Goat anti-rabbit HRP-conjugated secondary antibody (SouthernBiotech 4030-05), SuperSignal West Pico ECL (Thermo Scientific 34580), and a BioRad ChemiDoc Touch were used to visual immunoblot signal.

### X-gal Staining

Tissues to be used for *in situ* X-gal staining were dissected and placed in cold PBS on ice until all tissues to be stained were removed from the mouse. The intestine, stomach, and bladder contents were removed, and the organs rinsed with ice cold PBS. Liver and kidney samples were cut along the coronal and sagittal axes and the pieces rinsed with cold PBS to remove as much blood as possible prior to fixation and staining. Brain samples were cut into 2-3 mm thick slices along both the coronal and sagittal axes. Mammary tissue and skin samples were spread onto mesh biopsy bags to maintain orientation during fixation.

After harvest, tissues were fixed for 1 hour at 4°C in 4% formaldehyde solution (from a 37% stock freshly diluted in PBS). Tissues were washed once with X-gal rinse buffer (100 mM sodium phosphate pH 7.3, 2 mM MgCl_2_, 0.01% sodium deoxycholate, 0.02% NP-40) for 1 hour at 4°C, and two additional times for 30 minutes each at room temperature. All fixation and washing steps were performed with gentle agitation on an orbital rocker. Samples were then incubated in X-gal staining solution (X-gal rinse buffer plus 1 mg/mL X-gal (GoldBio X4281C, diluted from a 25 mg/mL stock in dimethylformamide), 1 mM potassium ferricyanide, 1 mM potassium ferrocyanide) for 48 hours at 37°C, after which they were post-fixed overnight in 10% NBF and placed in 70% reagent alcohol. Samples were photographed and then processed for embedding and sectioning.

### Clearing X-gal stained whole mounts

Tissues stained with X-gal as above were processed to increase optical clarity for brightfield photography (Schatz et al., 2005), with minor modifications. Briefly, after completion of X-gal staining, tissues were incubated in solutions of increasing glycerol concentrations (20%, 50%, 80% and 100% v/v), brought to final volume in 1% KOH (w/v) and incubated at 37°C for 2 days in each glycerol solution. Imaging was performed in 100% glycerol.

### Immunofluorescence

FFPE tissues were sectioned at 4-6 microns, baked at 60°C for 60 minutes prior to deparaffinization with xylene and rehydration through graded alcohols. Antigen retrieval was performed with pH 6.0 citrate buffer antigen retrieval solution (Sigma #C9999) in a pressure cooker and cooled for 20 minutes at room temperature. Sections were blocked with 10% normal goat serum in PBS+0.5% Tween, followed by incubation with rabbit-anti β-galactosidase (Invitrogen #A11132) and goat anti-rabbit IgG AlexaFluor488 (Invitrogen #A11008), both in Dako Antibody Diluent (Dako #S0809). TrueVIEW Autofluorescence Quenching reagent (Vector Laboratories #SP-8500) was applied to each section prior to coverslipping with Vectashield containing DAPI. Slides were imaged within 48hr after staining.

### Evaluating litter success

Litter success data were compiled retrospectively from colony breeding records. Litter records were evaluated for pup date of birth, entries for pup and/or litter loss, and the number of pups successfully weaned at 21 days of age. Litter success was defined as having at least one pup reaching weaning age.

## Results

We obtained mice carrying the ‘knockout-first’ *Grb7* tm1a *lacZ* gene trap allele from EUCOMM (Skarnes et al., 2011) and successive crosses with mice with ubiquitous FLP and Cre recombinase expression were used to generate mice bearing *Grb7*-null alleles (Fig 1A). PCR of tail-biopsy derived genomic DNA revealed successful genetic recombination events (Fig 1b). *Grb7*^*-/-*^ mice are viable to adulthood and are fertile. Elimination of Grb7 protein expression (and replacement by beta-galactosidase expression in the *lacZ* gene trap) was confirmed by western blot of liver lysates (Fig 1C).

Initial observations revealed no overt physical or behavioral abnormalities, so we submitted these mice for a systematic phenotyping necropsy by the University of Wisconsin Comparative Pathology Lab for analysis of all organ systems. This also revealed no gross or histopathological abnormalities (results summarized in Supplemental Table 1). To thoroughly examine tissues and cellular compartments in which *Grb7* is normally expressed for potential impacts of Grb7 elimination, we used the *Grb7*^*lacZ/lacZ*^ line (Fig 1A) in which the *lacZ* reporter replaces Grb7 expression. Because non-specific X-gal hydrolysis can occur in some tissues and to avoid issues caused by incomplete X-gal perfusion in solid organs (Sanchez-Ramos et al., 2000), we used immunofluorescence of tissue sections to visualize β-galactosidase activity.

In the brain, several regions exhibit reporter expression such as the pyramidal neurons in the cerebellar cortex (Fig 2A, B), the pyramidal cells and the inner granular layer of the cerebral cortex (Fig 2C, D, E), the hypothalamus (not shown) and the choroid plexus (Fig 2F). The eye shows reporter activity in the sclera and ciliary body (Fig 2G). Specifically, the retina exhibits reporter activity in the optic disc region (Fig 2H), ganglion cell layer, inner plexiform layer, outer plexiform layer, and the photoreceptor layer (Fig 2I). Reporter is detected in both white and gray matter of the spinal cord (Fig 2J), with enrichment in the dorsal and ventral horns.

**Figure 2.**
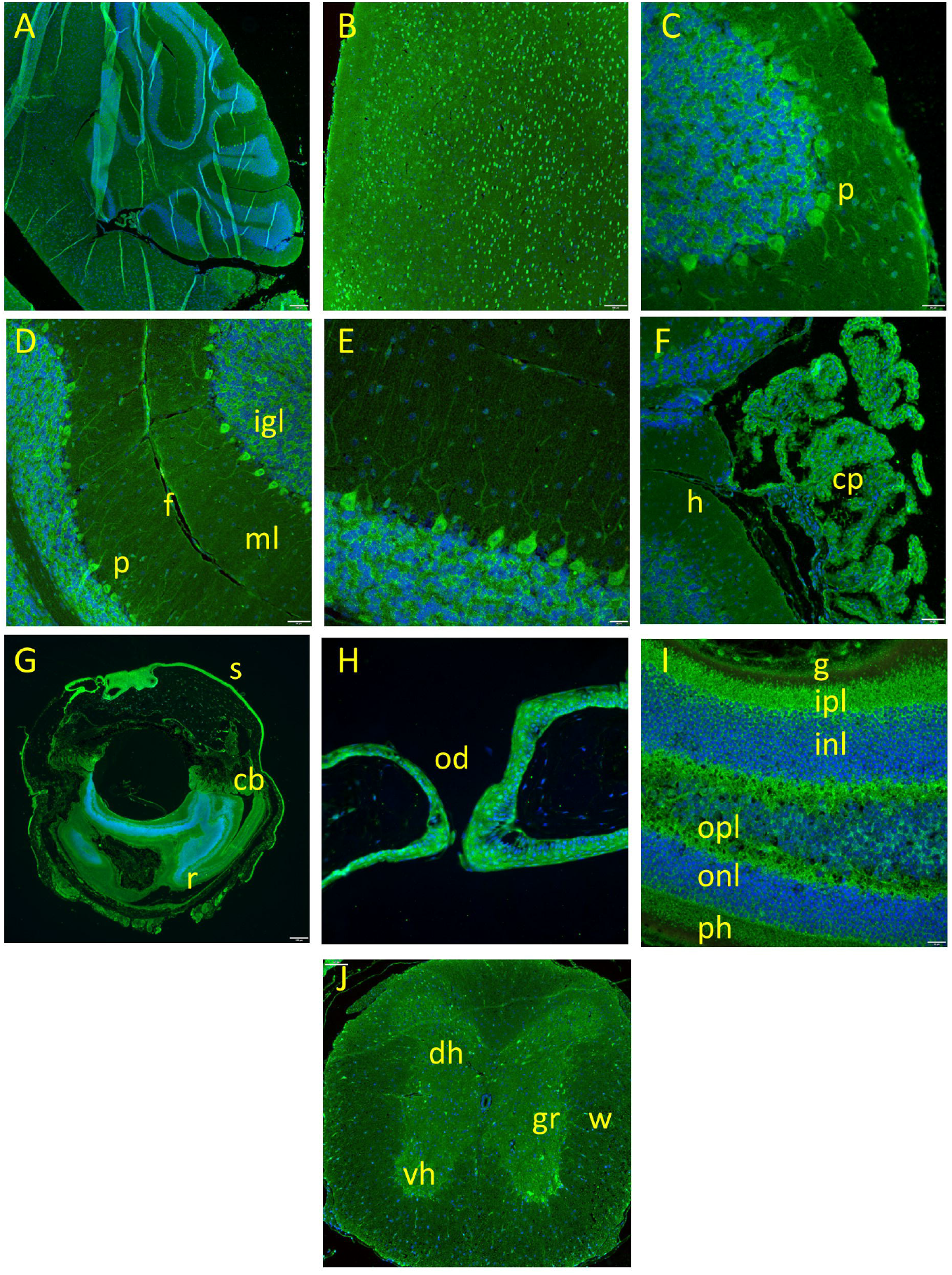
*Grb7* reporter localization in nervous system tissues. Anti β-galactosidase immunofluorescence (green) and nuclear counterstain (DAPI, blue) in *Grb7*^*lacZ/lacZ*^ mice. **A**. Cerebellar cortex, 4x. **B**. Cerebellar cortex, 10x. **C**. Cerebral cortex, 40x. (p) Purkinje cells **D**. Cerebral cortex, 10x. (igl) inner granular layer, (ml) molecular layer, (f) fissure, (p) Purkinje cells **E**. Cerebral cortex, 40x. **F**. Brain, 20x. (cp) choroid plexus, (h) hippocampus. **G**. Eye, 4x. (s) sclera, (cb) ciliary body, (r) retina. **H**. Eye, 40x. (od) optic disc. **I**. Retina, 40x. (g) granular layer, (ipl) inner plexiform layer, (inl) inner nuclear layer, (opl) outer plexiform layer, (onl) outer nuclear layer, (ph) photoreceptor layer. **J**. Spinal cord, cervico-thoracic region, 4x. (dh) dorsal horn, (vh) ventral horn, (gr) gray matter, (w) white matter.

Respiratory tissues of the tracheal epithelium (Fig 3A) are positive for β-galactosidase expression, as are the lungs (Fig 3B), primarily in the bronchioles and bronchiolar ducts (Fig 3C). Reporter expression was detected in the squamous epithelium of the esophagus (Fig 3D), the glandular and squamous epithelium of the stomach (Fig 3E), and the columnar epithelium of the small intestine crypts and submucosal glands (Fig 3F). Expression was also detected in the tubular epithelium of the kidney (Fig 3G), and the urothelial cells of the urinary bladder (Fig 3H) and the ureter (Fig 3I). Additionally, hepatocytes throughout the liver (Fig 3J) and the secretory epithelium of the gall bladder (Fig 3K) were strongly positive.

**Figure 3.**
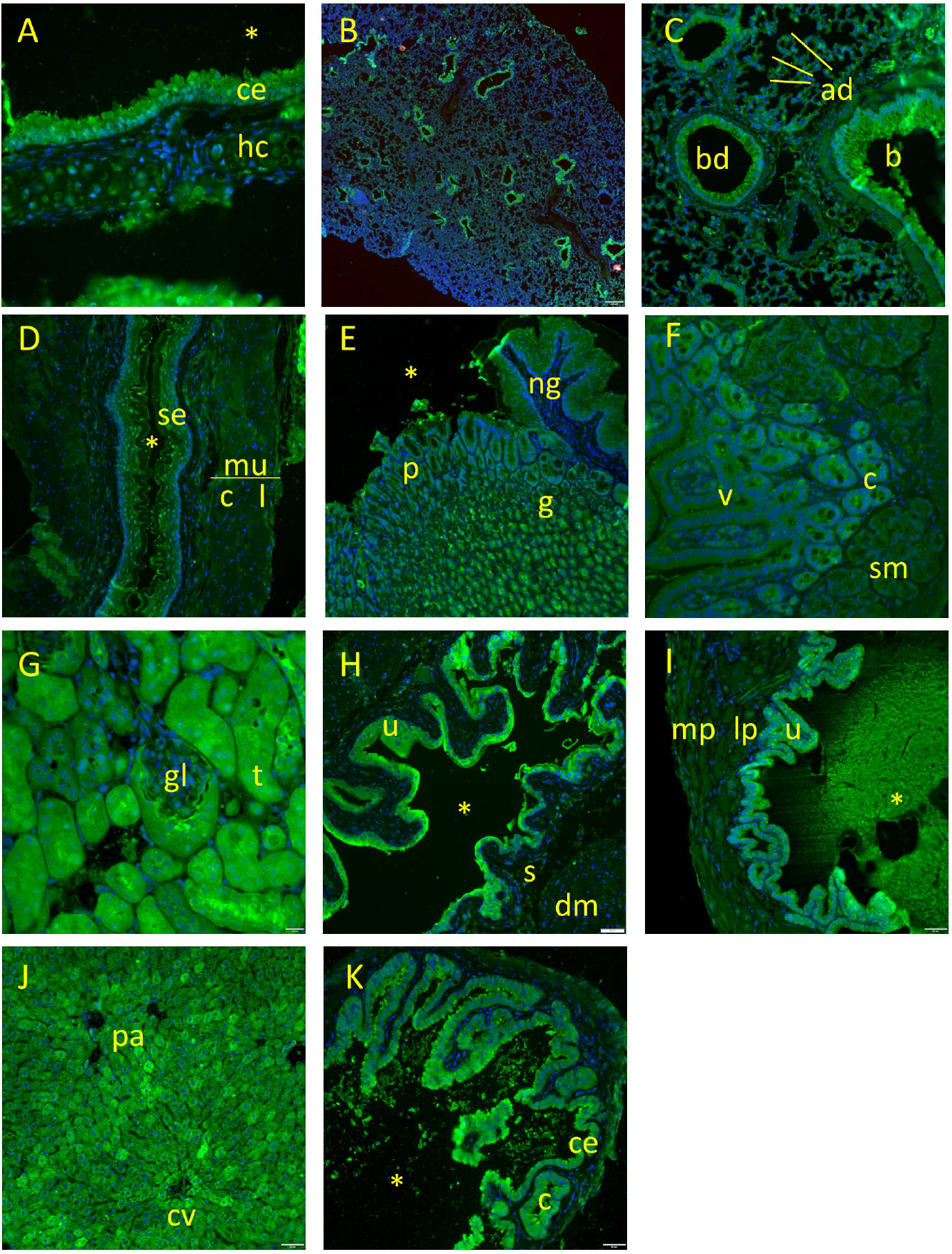
*Grb7* reporter localization in respiratory, digestive, and excretory tissues. Anti β-galactosidase immunofluorescence (green) and DAPI (blue) in *Grb7*^*lacZ/lacZ*^ mice. **A**. Trachea, 40x. (*) lumen, (ce) columnar epithelium, (hc) hyaline cartilage. **B**. Lung, 4x. **C**. Lung, 20x. (b) bronchiole, (bd) bronchiolar duct, (ad) alveolar duct. **D**. Esophagus, 20x. (*) lumen, (se) stratified squamous epithelium, (mu) muscular layer: (c) circular layer, (l) longitudinal layer. **E**. Stomach, 10x objective capture, enlarged in figure. (*) lumen, (ng) non-glandular stomach, (g) glandular stomach, (p) gastric pits. **F**. Small intestine, 10x objective capture, enlarged in figure. (v) villi, (c) crypt, (sm) submucosal gland. **G**. Kidney, 40x. (gl) glomerulus, (t) tubule. **H**. Urinary bladder, 20x. (*) lumen, (u) urothelium, (s) submucosa, (dm) detrusor muscle. **I**. Ureter, 20x. (mp) muscularis propria, (lp) lamina propria, (u) urothelium, (*) lumen. **J**. Liver, 20x. (pa) portal area, (cv) central vein. **K**. Gallbladder, 20x. (*) lumen, (ce) columnar epithelium, (c) crypt.

The ductal and glandular compartments of the salivary glands (Fig 4A), pancreatic glands and islets (Fig 4B), as well as the glandular thyroid follicles (Fig 4C) were positive for reporter. The pituitary (Fig 4D-F) exhibits intense staining in the anterior pituitary with a heterogeneous staining pattern at the cellular level. There was a clear absence of staining in the posterior pituitary. In the skin, sebaceous glands and portions of the hair follicles express β-galactosidase (Fig 4G-I).

**Figure 4.**
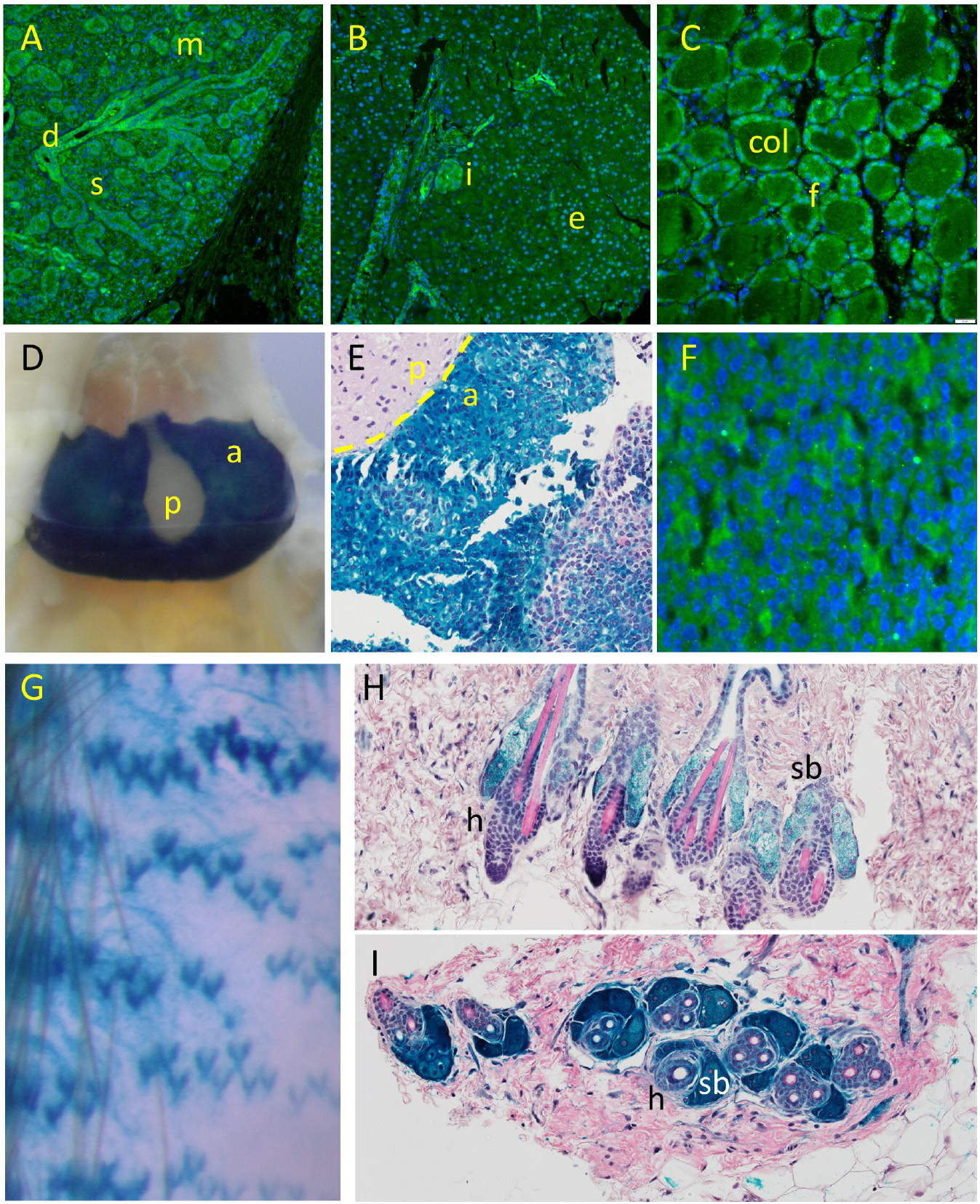
*Grb7* reporter localization in secretory and endocrine tissues. **A**. Salivary gland, 20x objective. (m) mucinous acini, (s) serous acini, (d) duct. **B**. Pancreas, 20x. (e) exocrine pancreas, (I) islet of Langerhans. **C**. Thyroid, 40x. (f) follicle, (col) colloid. **D**. Pituitary, in situ X-gal stain, whole mount. (a) anterior, (p) posterior. **E**. Pituitary, in situ X-gal stain followed by H&E stain, 20x capture, magnified in figure. (a) anterior, (p) posterior. **F**. Anterior pituitary, β-galactosidase immunofluorescence, 20x capture, magnified in figure. **G**. Skin, in situ X-gal stain, glycerol cleared whole mount. **H**. Skin, longitudinal section, in situ X-gal stain with H&E, 20x. (h) hair follicle, (sb) sebaceous gland. **I**. Skin, transverse section, in situ X-gal stain with H&E, 20x. (h) hair follicle, (sb) sebaceous gland. Immunofluorescence: anti β-galactosidase immunofluorescence (green), DAPI (blue). In situ X-gal (blue), with hematoxylin (purple) and eosin (pink/red).

Reporter-positive female reproductive structures (Fig 5A) include the oviduct (Fig 5A), the corpus luteum in the ovary (Fig 5 A,B), glandular and luminal uterine tissue (Fig 5A,C), and placenta (not shown). Male reproductive tissues (Fig 5D) showed reporter presence in the testis (Fig 5D,E), epididymis (Fig 5D,F), preputial gland (Fig 5G), vas deferens (Fig 5 D,H), and the prostate (Fig 5I).

**Figure 5.**
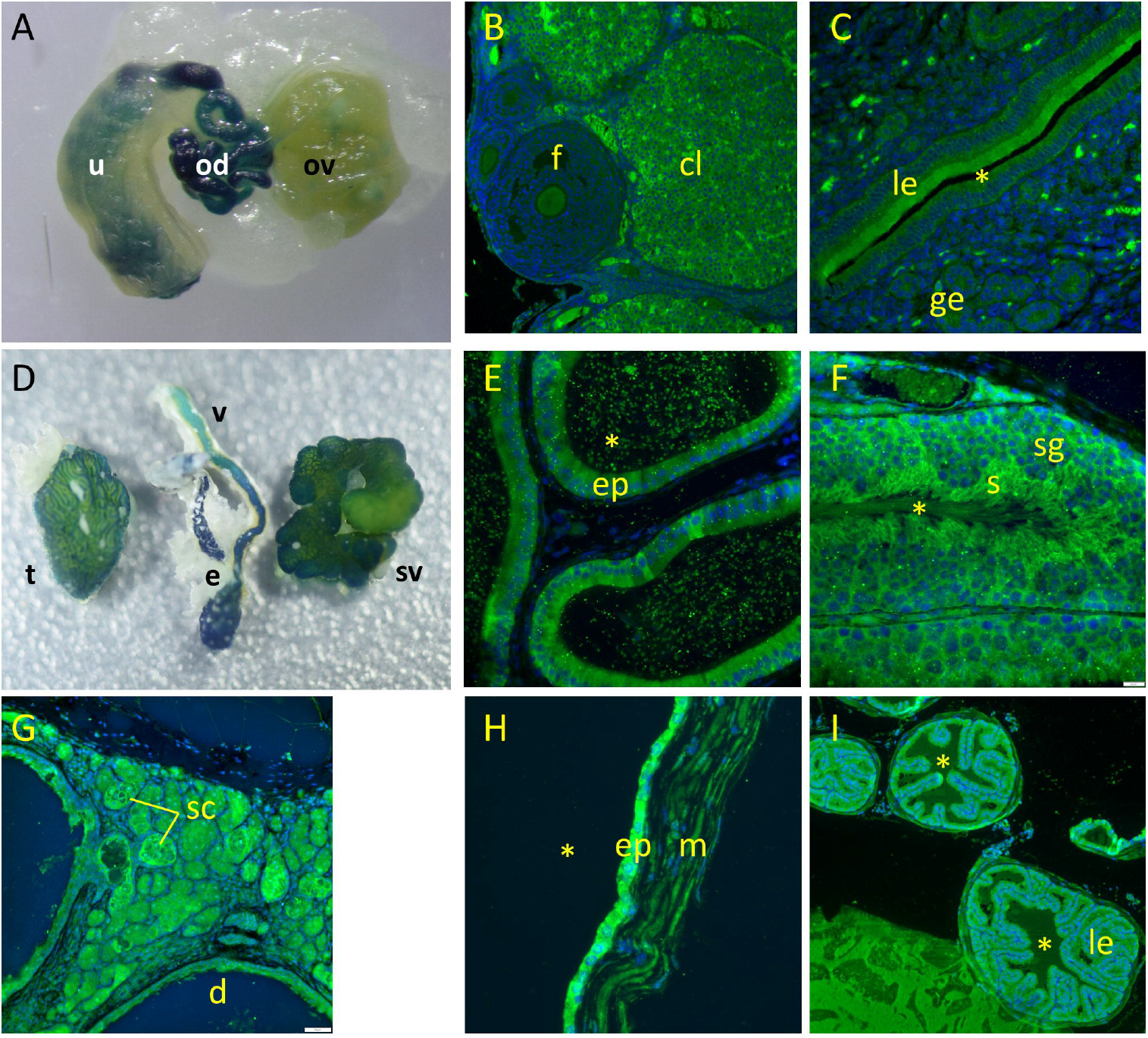
*Grb7* reporter localization in reproductive tissues. **A**. Female reproductive organs, in situ X-gal whole mount. (ov) ovary, (od) oviduct, (u) uterus, partial. **B**. Ovary, 20x. (f) follicle, (cl) corpus luteum. **C**. Uterus, 40x. (*) lumen, (le) lumenal epithelium, (ge) glandular epithelium. **D**. Male reproductive organs, in situ X-gal whole mount. (t) testis, (e) epididymis, (v) vas deferens, (sv) seminal vesicle. **E**. Epididymis, 40x. (*) lumen, (ep) epithelium. **F**. Testis, 40x. (*) lumen, (sg) spermatogenic epithelium, (s) sperm. **G**. Preputial gland, 20x. (sc) secretory cells, (d) duct. **H**. Seminal vesicle, 40x. (*) lumen, (ep) epithelium, (m) muscular layer. **I**. Prostate gland, 20x. (*) lumen, (le) lumenal epithelium. Immunofluorescence: anti β-galactosidase (green), DAPI (blue). In situ X-gal whole mount (blue).

The mammary gland fully arborized the fat pad (Fig 6A) and exhibited reporter activity in several compartments of the structure. In situ X-gal staining revealed the entire ductal tree was positive for reporter (Fig 6A, whole mount mammary gland and 6B, nipple and collecting duct in a glycerol clarified whole mount). Specific regions of the nipple were positive (Fig 6C, D) and the collecting duct was clearly positive and distinguishable from the epidermis of the nipple (arrowheads in Fig 6C, D). Histological analysis confirmed the circumferential reporter localization around the perimeter of the nipple (Fig 6E) and the luminal epithelium in the mammary duct (Fig 6G). Reporter expression was also evident in the alveolar buds.

**Figure 6.**
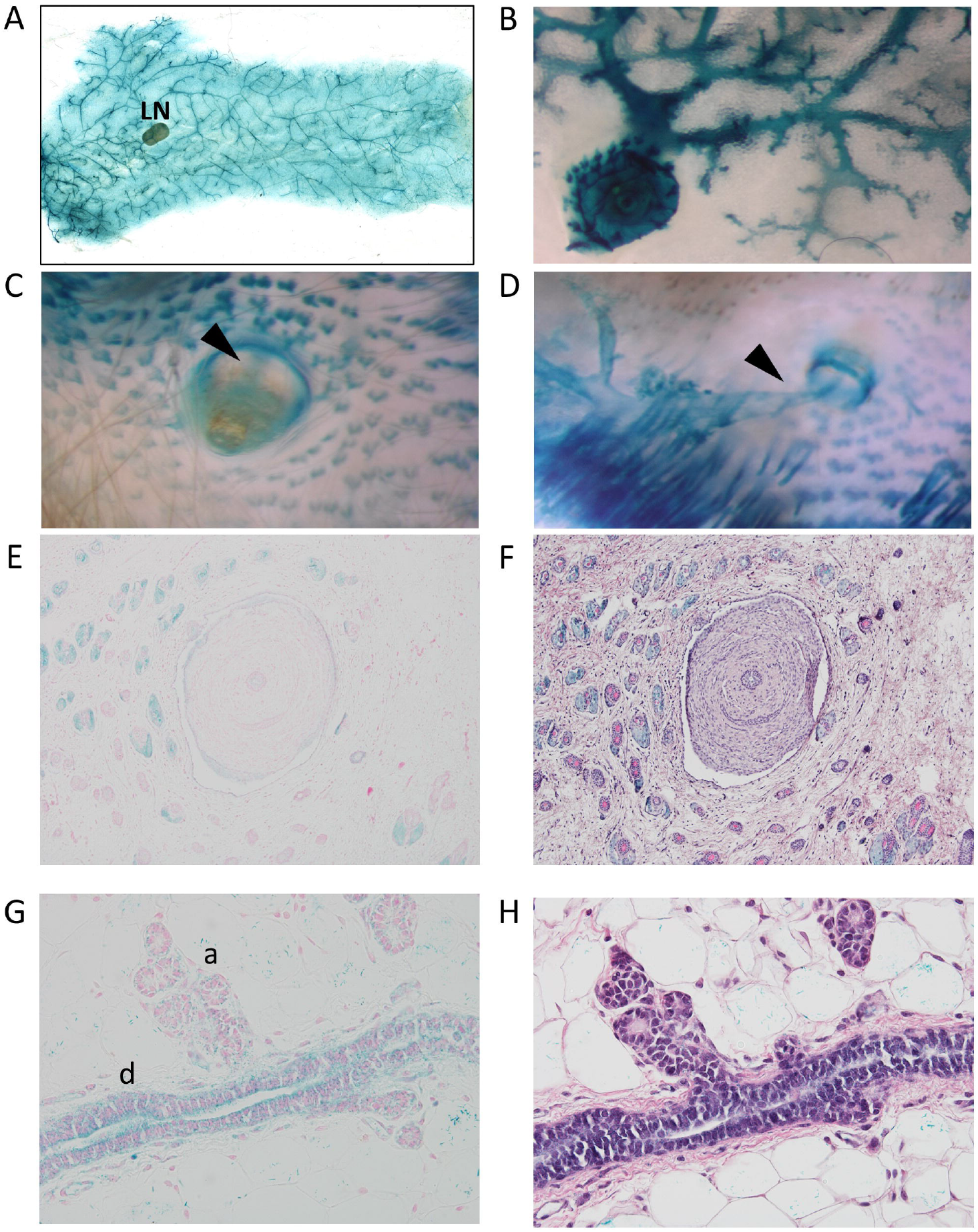
Mammary *Grb7* reporter localization. **A**. Mammary gland, in situ X-gal stained whole mount. (LN) lymph node. **B**. Nipple and collecting duct, X-gal stained mammary gland whole mount, magnified region. **C**. Nipple, skin, and collecting mammary duct, external to internal view. Arrowhead: mammary duct. **D**. Nipple, skin, and collecting mammary duct, internal to external view. Arrowhead: mammary duct. **E**. Nipple, histological section from tissue after in situ X-gal staining and nuclear fast red counterstain (red/pink), 10x. **F**. H&E stained serial section of E, 10x. **G**. Mammary duct, histological section from tissue after in situ X-gal staining and nuclear fast red counterstain (red/pink), 20x. **H**. H&E stained serial section of G, 20x.

Although *Grb7* knockout mice seem to develop quite normally, we experienced significant challenges in colony management. *Grb7* heterozygous and knockout females do not experience any difficulty becoming pregnant but had a striking dissimilarity in their ability to successfully raise litters to weaning age. 70% of litters born to *Grb7* heterozygous females were successfully raised (at least one pup weaned at 21 days), while only 9% of litters born to *Grb7* knockout females succeeded (Fisher’s Exact Test P < 0.0001, Fig 7A). This failure was most commonly exemplified by death of most or all of a litter in the early post-natal period (days 0-2). To determine if paternal genotype had an impact, we specifically examined litter success in *Grb7* heterozygous females crossed with either *Grb7* heterozygous or knockout males. There was no significant difference in litter success associated with paternal genotype (Fisher’s Exact Test P = 0.26). In addition, we noted no survival disparity between pups in litters born to *Grb7* heterozygous males and *Grb7* knockout females (in which the expected Mendelian ratio of Grb7 heterozygous pups is 0.5). All pups were impacted in cases of litter demise. This observed pattern is most consistent with a maternal failure to provide enough milk to support pup growth.

**Figure 7.**
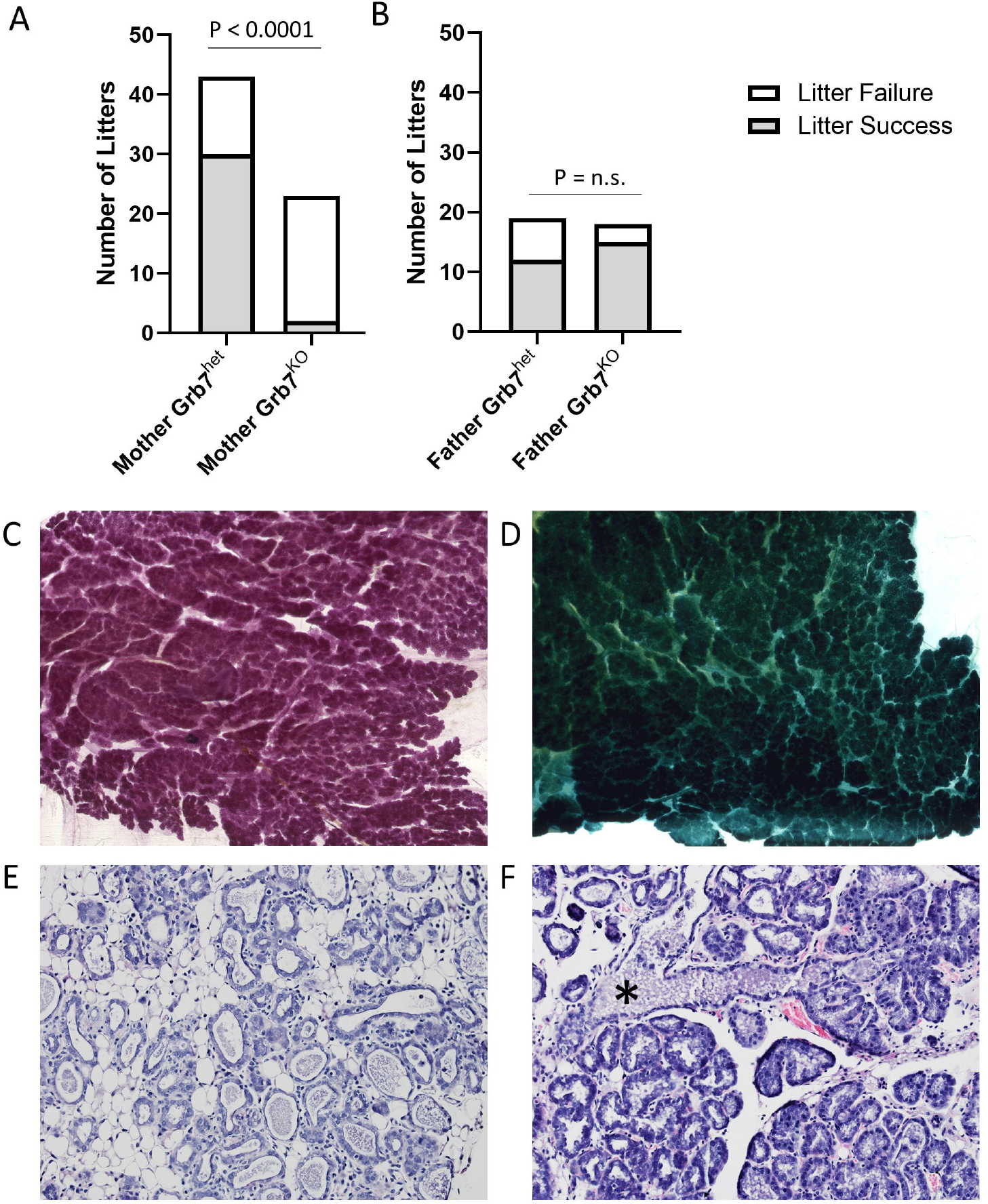
Pup demise and functional differentiation of the mammary gland. **A**. Litter success stratified by maternal genotype. **B**. Litter success in Grb7 heterozygous females stratified by paternal genotype. **C**. Inguinal mammary gland whole mount, carmine stain, from a lactating *Grb7*^*lacZ/lacZ*^ female, 7 days post-partum. **D**. Thoracic mammary gland whole mount, in situ X-gal stain, from a lactating *Grb7*^*lacZ/lacZ*^ female, 7 days post-partum. **E**. *Grb7*^*-/-*^ mammary gland, 20x. Estimated pregnancy day 16. **F**. *Grb7*^*-/-*^ mammary gland, 20x. lactation day 11, showing milk collection in the duct (*).

Functional differentiation of the mammary gland appeared normal by whole mount analysis. Carmine alum staining illustrates that expansion of the epithelial compartment occurs with pregnancy (Fig 7C) and in situ X-gal staining indicates that the expanding population expresses β-galactosidase reporter (Fig 7D). Histological analysis of mammary glands from pregnant (Fig 7E) and lactating (Fig 7F) females shows milk production, with progression into the duct (asterisk, Fig 7F). Accordingly, *Grb7* knockout mammary epithelial cells appear capable of synthesizing and appropriately secreting milk into mammary ducts, although neither the nutritional content of the milk nor the efficiency of milk ejection was evaluated here.

Additional tissues assayed that lacked reporter activity include the lymphohematopoietic tissues of the spleen, thymus, lymph nodes, and bone marrow and the main musculoskeletal system including cardiac, skeletal (hind limb), and smooth muscle (bladder) samples. Bone samples taken from the rib and femur were also negative.

## Discussion

Given the breadth of signaling pathways that Grb7 participates in as an adapter protein, we hypothesized that generating a *Grb7*-null mouse would reveal critical roles for Grb7 in mouse development, organogenesis, and/or physiology. To our surprise, despite clearly eliminating Grb7 expression (Fig 1), the whole-body knockout mouse that we derived did not have an overt phenotype at gross or histological levels of analysis (Supplemental Table 1). Using *lacZ* expression as a marker for cell lineages in which the *Grb7* promoter is active also did not reveal obvious architectural defects.

Normal expression of *Grb7* in the mouse (Figs 2-6) spans multiple organ systems, predominantly localized to the epithelial compartment of tissues and, in particular, glandular epithelium. Additionally, the reporter-tagged *Grb7* allele was active in some cell lineages of the central nervous system (in brain, eye, and spinal cord). The broad expression pattern implies that Grb7 may have roles in multiple tissues, but the lack of an observed developmental phenotype in the knockout suggests that there may be significant redundancy built into these pathways, with Grb10 and/or Grb14 potentially able to rescue any defects resulting from Grb7 loss.

The only striking phenotype observed in these animals was the early post-natal death of pups born to *Grb7* knockout females, which impacted both *Grb7* knockout and heterozygous pups. Grb7 is expressed in the mammary gland (Fig 6) and the glands in *Grb7* knockout mice appeared quite normal. Potential explanations include (1) a mammary-specific failure to synthesize one or more key nutritional components of milk, (2) a lactational failure due to a mammary-specific requirement for Grb7 in either milk secretion or mammary gland contractility, (3) a failure to maintain lactation after initiation (i.e. an early involution) or (4) a distant impact caused by a requirement for Grb7 for the function of an endocrine gland (e.g. pituitary or ovary) that has a key role in regulating mammary gland physiology and function.

While Grb7 appears dispensable for the many aspects of development examined here, we have not addressed its potential role in cancer, in which *Grb7* amplification and overexpression has been reported in several tumor types. By breeding with mouse models which develop tumors in these organs, it may be possible to determine the specific contribution, if any, of Grb7 to tumorigenesis, metastasis or therapy response.

The deregulation of GRB7 in multiple tumor tissues has prompted the development of several candidate inhibitors (Ambaye et al., 2011; Pero et al., 2002; Watson et al., 2017) with activity in preclinical models. The cyclic peptide inhibitor, G718NATE, had demonstrable activity against xenografted human cancer cell lines and had no obvious toxicity in mice (Tanaka et al., 2006). Our demonstration here that Grb7 is dispensable for mouse development and most aspects of adult homeostasis examined supports the idea that GRB7 inhibition in adult humans would be tolerable.

In conclusion, the relatively minor phenotype in *Grb7*-deleted mice was surprising to us, given the extensive prior reports on the involvement of Grb7 in so many signaling pathways (Chu et al., 2019; Han et al., 2001). Intercrossing with mice deficient in other, potentially compensating, Grb gene family members may be necessary to fully elucidate the key role(s) of this family of adapter proteins in mammalian physiology.

## Competing interests

The authors declare that they have no competing interests.

## Acknowledgements

This study was funded by the Gundersen Medical Foundation. PK holds the Dr. Jon & Betty Kabara Endowed Chair in Precision Oncology. KL was supported by the Norman L. Gillette, Jr. Cancer Research Fellowship.

## Authors’ contributions

KL and PK both devised and performed experiments, analyzed the data, and wrote the manuscript. Both authors approved the final manuscript.

## Notes

### Competing Interest Statement

The authors have declared no competing interest.

