## Supplemental Table 1 for "Grb7 knockout mice develop normally but litters born to knockout females fail to thrive"

Supplemental Table 1: Grb7 Knockout Mouse Phenotyping - Necropsy Observations

| **System** | **Organ/tissues** | **Gross Finding** | **Histopathology (H&E)** |
| --- | --- | --- | --- |
| Integument | Coat, skin, subcutaneous fat  Mammary Gland | Unremarkable  Not described | Unremarkable  Unremarkable |
| Cardiovascular | Heart | Unremarkable | Unremarkable |
| Respiratory | Trachea  Lungs  Nasal Cavity | Unremarkable  Diffuse, coral pink  Not described | Unremarkable  Open and clear alveolar spaces, airways  Unremarkable epithelium and stroma |
| Digestive | Oral cavity, teeth, tongue  Salivary Glands  Esophagus  Stomach  Pancreas  Sm. Intestine, cecum, colon  Liver | Unremarkable  Not described  Not described  Contains ingesta  Pale pink, unremarkable  Contain digesta, feces  Pale brown, sharp edges | Unremarkable  Unremarkable  Moderate hyperkeratosis on squamous epithelium  Mild hyperkeratosis on squamous epithelium; normal glandular  Unremarkable  Unremarkable  Unremarkable, moderate variation in nuclear size |
| Lymphohematopoietic | Spleen  Thymus  Bone Marrow (femur) | Deep red/brown  White/grey, normal volume  Dark red, no obvious abnormality | Large lymphoid follicles, mild mantle zone hyperplasia  Not performed  Unremarkable, myeloid: erythroid ratio adequate |
| Urogenital/Reproductive | Kidneys  Bladder  Testes, Seminal vesicles, Prostate  Uterus  Ovaries | Unremarkable  Unremarkable  Unremarkable in size and maturity  Not described  Mild mucometra, moderate fat stores  Not described | Adequate no. of small glomeruli, normal proximal/distal tubules  Not performed  Unremarkable, normal spermatogenesis, gland, secretion  Unremarkable epithelium and stroma  Defined uterine glands (estrus)  Normal/post-ovulatory follicles present |
| Endocrine | Adrenal  Thyroid, Parathyroid | Apparent normal color, size  Apparent normal color, size | Not performed  Unremarkable, variably sized follicles w eosinophilic colloid |
| CNS | Brain  Eyes | No gross external abnormalities  Unremarkable | Moderate nuclear vacuolation  Unremarkable |
| Musculoskeletal | Muscular  Skeletal | No gross lesions or abnormalities  No gross lesions or abnormalities | Unremarkable  Unremarkable bony sections |
